# GrandPrix: Scaling up the Bayesian GPLVM for single-cell data

**DOI:** 10.1101/227843

**Authors:** Sumon Ahmed, Magnus Rattray, Alexis Boukouvalas

## Abstract

**Motivation:** The Gaussian Process Latent Variable Model (GPLVM) is a popular approach for dimensionality reduction of single-cell data and has been used for pseudotime estimation with capture time information. However current implementations are computationally intensive and will not scale up to modern droplet-based single-cell datasets which routinely profile many tens of thousands of cells.

**Results:** We provide an efficient implementation which allows scaling up this approach to modern single-cell datasets. We also generalize the application of pseudotime inference to cases where there are other sources of variation, such as branching dynamics. We apply our method on microarray, nCounter, RNA-seq, qPCR and droplet-based datasets from different organisms. The model converges an order of magnitude faster compared to existing methods whilst achieving similar levels of estimation accuracy. Further, we demonstrate the flexibility of our approach by extending the model to higher-dimensional latent spaces that can be used to simultaneously infer pseudotime and other structure such as branching. Thus, the model has the capability of producing meaningful biological insights about cell ordering as well as cell fate regulation.

**Availability:** Software available at github.com/ManchesterBioinference/GrandPrix.

## 1 Introduction

The analysis of single-cell genomics data promises to reveal novel states of complex biological processes, but is challenging due to inherent biological and technical noise. It is often useful to reduce high-dimensional single-cell gene expression profiles into a low-dimensional latent space capturing major sources of inter-cell variation in the data. Popular methods for dimensionality reduction applied to single-cell data include linear methods such as Principal and Independent Components Analysis (P/ICA) (Trapnell *et al*., 2014; Ji and Ji, 2016) and nonlinear techniques such as t-stochastic neighbourhood embedding (t-SNE) (Becher *et al*., 2014), diffusion maps (Haghverdi *et al*., 2015, 2016) and the Gaussian Process Latent Variable Model (GPLVM) (Buettner and Theis, 2012; Buettner *et al*., 2015). In some cases the dimension is reduced to a single *pseudotime* dimension representing the trajectory of cells undergoing some dynamic process such as differentiation or cell division. The pseudotemporal ordering of cells is based on the principle that cells represent a time series where each cell corresponds to distinct time points along the pseudotime trajectory, corresponding to progress through a process of interest. The trajectory may be linear or branching depending on the underlying process.

Different formalisms can be used to represent a pseudotime trajectory. In graph-based methods such as Monocle (Trapnell *et al*., 2014), Wanderlust (Bendall *et al*., 2014), Waterfall (Shin *et al*., 2015) and TSCAN (Ji and Ji, 2016), a simplified graph or tree is estimated. By using different path-finding algorithms, these methods try to find a path through a series of nodes. These nodes can correspond to individual cells (Trapnell *et al*., 2014; Bendall *et al*., 2014) or groups of cells (Shin *et al*., 2015; Ji and Ji, 2016) in the graph. SCUBA (Marco *et al*., 2014) uses curve fitting to characterize the pseudotime trajectory. Principal curves are used to model the trajectory and each cell is assigned a pseudotime according to its low-dimensional projection on the principal curves. On the other hand, in the diffusion pseudotime (DPT) framework (Haghverdi *et al*., 2016), there is no initial dimension reduction. DPT uses random walk based inference where all the diffusion components are used to infer pseudotime.

One major drawback of the above methods is the absence of an explicit probabilistic framework. They only provide a single point estimate of pseudotime, concealing the impact of biological and technical variability. Thus, the inherent uncertainty associated with pseudotime estimation is not propagated to the downstream analysis and its consequences remain unknown. However, the robustness of the estimated pseudotime for these models can be examined by re-estimating the pseudotimes multiple times under different initial conditions, parameter settings or samples of the original data. Campbell and Yau (2016) have examined the pseudotime estimation of Monocle where they have taken multiple random subsets of data and re-estimated the pseudotimes for each of them. They have shown that the pseudotime points assigned by Monocle for the same cell can vary significantly across the random subsets taken. This uncertainty in pseudotime assignment motivates the use of probabilistic analysis techniques. The GPLVM is a non-linear probabilistic model for dimension reduction (Lawrence, 2005) and has been used extensively to analyse singlecell data. Buettner and Theis (2012) used the GPLVM for non-linear dimension reduction to uncover the complex interactions among differentiating cells. Buettner *et al*. (2015) used the GPLVM to identify subpopulations of cells where the algorithm also dealt with confounding factors such as cell cycle. More recently, Bayesian versions of the GPLVM have been used to model pseudotime uncertainty. Campbell and Yau (2016) have proposed a method using the GPLVM to model pseudotime trajectories as latent variables. They used Markov Chain Monte Carlo (MCMC) to draw samples from the posterior pseudotime distribution, where each sample corresponds to one possible pseudotime ordering for the cells with associated uncertainties. Zwiessele and Lawrence (2016) have used the Bayesian GPLVM framework to estimate the Waddington landscape using single-cell transcriptomic data; the probabilistic nature of the model allows for more robust estimation of the topology of the estimated epigenetic landscape.

As well as allowing for uncertainty in inferences, Bayesian methods have the advantage of allowing the incorporation of additional covariates which can inform useful dimensionality reduction through the prior. In particular, pseudotime estimation methods may usefully incorporate capture times which may be available from a singlecell time series experiment. For example, in the immune response after infection, gene expression profiles show a cyclic behaviour which makes it challenging to estimate a single pseudotime. Reid and Wernisch (2016) have developed a Bayesian approach that uses a GPLVM with a prior structure on the latent dimension. The latent dimension in their model is a one-dimensional pseudotime and the prior relates it to the cell capture time. This helps to identify specific features of interest such as cyclic behaviour of cell cycle data. The pseudotime points estimated by their model are in proximity to the actual capture time and use the same scale. Further, Lönnberg *et al*. (2017) have adopted this approach and used sample capture time as prior information to infer pseudotime in the their trajectory analysis.

However, although the Bayesian GPLVM provides an appealing approach for pseudotime estimation with prior information, existing implementations are too computationally inefficient for application to large single-cell datasets, e.g. from droplet-based RNA-Seq experiments. In this contribution, we develop a new efficient implementation of the Bayesian GPLVM with an informative prior which allows for application to much larger datasets than previously considered. Furthermore, we show how extending the pseudotime model to include additional latent dimensions allows for improved pseudotime estimation in the case of branching dynamics. Our model is based on the variational sparse approximation of the Bayesian GPLVM (Titsias and Lawrence, 2010) that can generate a full posterior using only a small number of inducing points and is implemented within a flexible architecture (Matthews *et al*., 2017) that uses TensorFlow to perform computation across a number of CPU cores and GPUs.

## 2 Methods

Our model is motivated by the DeLorean approach (Reid and Wernisch, 2016) and uses cell capture time to specify a prior over the pseudotime. The probabilistic nature of the model can be used to quantify the uncertainty associated with pseudotime estimation. The GPLVM uses a Gaussian process (GP) to define the stochastic mapping from a latent pseudotime space to an observed gene expression space. A Gaussian process is an infinite dimensional multivariate normal distribution characterised by a mean function and a covariance function (Rasmussen and Williams, 2006). In the GPLVM, the mean function defines the expected mapping from the latent dimension to the observed data and the covariance function describes the associated covariance between the mapping function evaluated at any two arbitrary points in the latent space. Thus, the covariance function controls the second order statistics and can be chosen based on different second order features such as smoothness and periodicity.

### 2.1 Model

The challenge is to develop scalable models that can handle both biological and technical noise inherent in the data. Our preference for the sparse Bayesian approach offers a principled yet pragmatic answer to these challenges. The core of the model is the Gaussian process which has been used extensively to model uncertainty in regression, classification and dimension reduction tasks. The model uses a sparse variational approximation which requires only a small number of inducing points to efficiently produce a full posterior distribution.

The model we use is similar to the Bayesian GPLVM DeLorean model (Reid and Wernisch, 2016); the main differences between the two approaches lie in how model inference is accomplished which is discussed in Section 2.2. The primary latent variables in our method are the pseudotimes associated with each cell. The method expects the technical variability is sufficiently described by a Gaussian distribution which is often accomplished by taking a logarithmic transformation of the gene expression data. The critical assumption is that the cell capture times are informative for the biological dynamics of interest. The expression profile of each gene *y_g_* is modelled as a non-linear transformation of pseudotime which is corrupted by some noise *∊*

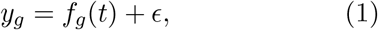

where 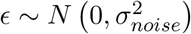 is a Gaussian distribution with variance 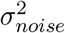. We place a Gaussian process prior on the mapping function

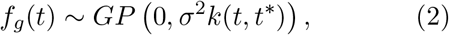

where *σ*^2^ is the process variance and *k*(*t, t*\*) is the covariance function between two distinct pseudotime points *t* and *t*\*. Thus, the expression profiles are functions of pseudotime and the covariance function imposes a smoothness constraint that is shared by all genes.

The pseudotime *t_c_* of cell *c* is given a normal prior distribution centred on the capture time *τ_c_* of cell *c*,

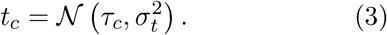

Here 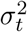 describes the prior variance of pseudotimes around each capture time.

To identify a non-periodic smooth pseudotime trajectory we have used the Radial Basis Function (RBF) and Matern_3/2_ kernels:

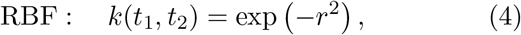

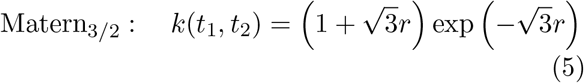

where 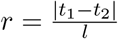 and *l* is the process length scale.

For cell cycle data, we have used the periodic kernel described in MacKay (1998). For a known period *λ*

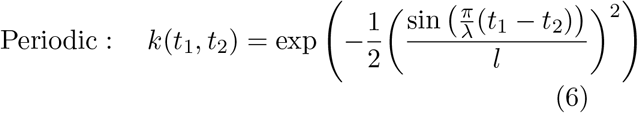

which limits the GP prior to periodic functions.

We have exploited the model’s flexibility by extending it to higher dimensional latent spaces. If the *x* represents the extra latent dimensions, then the expression profile of each gene is modelled as

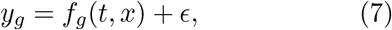

where

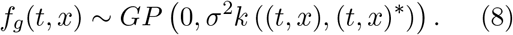

This generalisation takes the model beyond the estimation of pseudotime to provide a more general probabilistic non-linear dimension reduction technique.

### 2.2 Inference

The computation of the log marginal likelihood is mathematically intractable and MCMC methods (Campbell and Yau, 2016; Reid and Wernisch, 2016) have been employed for inference. Reid and Wernisch (2016) also use black box variational approaches that rely on data subsampling to increase inference efficiency. However, for the Bayesian GPLVM an analytic exact bound exists (Titsias and Lawrence, 2010; Damianou *et al*., 2016) but the original derivation and all currently available packages such as GPy (2012) assume an uninformative prior. We modify the exact bound to allow for informative priors

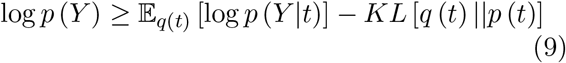

where *q* (*t*) is the variational distribution and

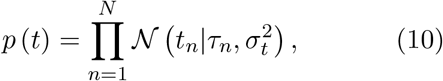

is the modified prior centred at the capture time *τ_n_* of cell *n* with prior variance 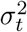. The variational approximation for the inputs *q* (*t*) is a factorised Gaussian as in the standard Bayesian GPLVM (Titsias and Lawrence, 2010)

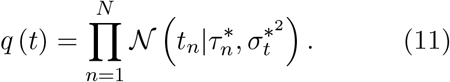

The modified lower bound on the model marginal likelihood is used to optimise all model parameters including the kernel hyperparameters (process variance, length scale, noise model variance) and the pseudotime locations. The Gaussian assumption for the variational approximate distribution may fail to adequately model multimodal distributions and model inference may be susceptible to local optima, as different pseudotime orderings may provide similarly smooth expression profiles. Careful initialisation of the mean 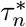 of variational approximation *q* (*t*) helps the algorithm to obtain good orderings (see Supplementary). Although using a non-Gaussian distribution would be possible, it would require a more complex approximate inference scheme (Rasmussen and Williams, 2006). In our experiments we find the estimated pseudotime ordering to be in close agreement with known times as reflected by high rank correlation values.

The most common practical limitation of GPs in practice is the computation required for inference; for each optimisation step the algorithm requires 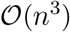 time and 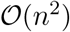 memory, where *n* is the number of training examples. Campbell and Yau (2016) have incorporated an MCMC implementation of the Bayesian GPLVM without an approximation in their model and hence their approach does not scale for large datasets.

The Bayesian GPLVM has computational complexity of 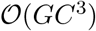, where *G* is the number of genes and *C* is the number of cells. To make the model computationally tractable for large datasets, a variety of sparse approximations have been proposed (Quiñonero-Candela and Rasmussen, 2005). Sparse GP approximations reduce the complexity to 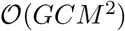 where *M* ≪ *C* is the number of auxiliary or inducing points. These inducing points may or may not coincide with actual points. As *M* is chosen much smaller than *C*, sparse approximations can result in significant reductions in computational requirements.

To reduce computational complexity Reid and Wernisch (2016) use the Fully Independent Training Conditional (FITC) approximation (Snelson and Ghahramani, 2006). This is a simple approach where a specific type of kernel is used to reduce the computational requirement. The approach is attractive because only the kernel is affected; the bound on the marginal likelihood is not affected and is therefore simple to derive. However as Bauer *et al*. (2016) have shown, this approach is prone to overfitting as it does not penalize model complexity. Titsias (2009) derived a Variational Free Energy (VFE) approximation for GP regression where the bound of the marginal likelihood is modified to include such a penalty term.

Both methods can be succinctly summarized by a different parametrisation of the marginal likelihood bound:

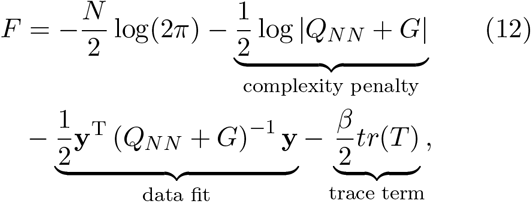

For the VFE approximation we have

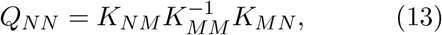

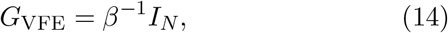

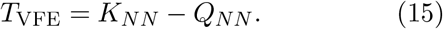

Here, *Q_NN_* is approximating the true covariance matrix *K_NN_*, but only involves the inversion of a *M* × *M* matrix *K_MM_. K_MM_* is the covariance matrix on the inducing inputs *Z*; *K_NM_* is the cross covariance matrix between the training and inducing inputs, i.e. between *X* and *Z* and 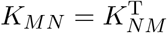.

The objective function of Equation (12) consists of three terms: the data fit term imposes a penalty on data not well explained by the model; the complexity term characterises the volume of probable datasets which are compatible with the data fit term and therefore penalises complex models fitting well on only a small ratio of datasets. Finally, the trace term measures the additional error due to the sparse approximation. Without this term VFE may overestimate the marginal likelihood like previous methods of sparse approximation such as FITC. In fact, the objective function of the FITC can be obtained from Equation (12) by using the same expression for *Q_NN_* and taking

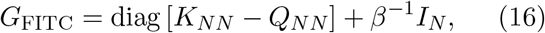

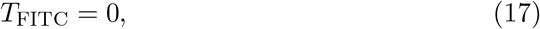

which clearly shows that the objective function of the FITC can be obtained by modifying the GP prior

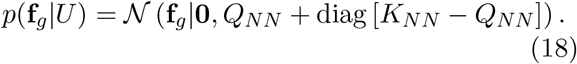

Here the inducing points are acting as an extra set of hyperparameters to parametrise the covariance matrix *Q_NN_*. As this approach changes the prior, the continuous optimisation of the latent variable **f**_*g*_ with respect to the inducing points *U* does not guarantee to approximate the full GP posterior (Titsias, 2009). Moreover, as **f**_*g*_ is heavily parametrised because of the extra hyperparameter *U* and the trace term is 0, overfitting may arise at the time of jointly estimating the inducing points and hyperparameters. For detailed derivation of the bound see the supplement and for a comprehensive comparison of FITC and VFE see Bauer *et al*. (2016). For both the VFE and FITC approximations, the inducing points may be chosen randomly from the training inputs or optimized with respect to the marginal likelihood bound.

Lastly, we have implemented our model in the GPflow package whose flexible architecture allows to perform the computation across multiple CPU cores and GPUs (Matthews *et al*., 2017).

The source of the scalability of our approach compared to DeLorean is therefore three-fold: model estimation using an exact variational bound, a robust sparse approximation (VFE vs FITC) and implementation on a scalable software architecture.

## 3 Results and discussion

The performance of our model has been investigated by applying it on a number of datasets of varying sizes collected from different organisms using different techniques. First we have compared our method with the DeLorean model (Reid and Wernisch, 2016) in terms of model fitting as well as the time required to fit the model on all the datasets used by Reid and Wernisch (2016); this encompasses the whole-leaf microarrays of *Arabidopsis thaliana* (Windram *et al*., 2012); single-cell RNA-Seq libraries of mouse dendritic cells (Shalek *et al*., 2014) and single-cell expression profiles of a human prostate cancer cell line (McDavid *et al*., 2014). Unlike the approach taken in Reid and Wernisch (2016) where the variational approximation is computed numerically, our approach provides an exact analytical bound which, as we show, results in robust parameter estimation. Moreover, our method converges quickly by using a small number of inducing points even for large data. Overall, our model outperforms the DeLorean model in both robustness and computational scalability aspects.

We also apply our approach on more recent droplet-based single-cell data. We apply the model on mouse embryo single-cell RNA-seq (Klein *et al*., 2015) and compare our the predicted pseudotime with results from the diffusion pseudotime method (DPT) (Haghverdi *et al*., 2016). We then apply the model on a large singlecell dataset of 3′ mRNA count data from peripheral blood mononuclear cells (Zheng *et al*., 2017) to demonstrate scalability to tens of thousands of cells.

Finally, we demonstrate the flexibility of the model by applying it on single-cell qPCR data of early development stages collected from mouse blastocyst (Guo *et al*., 2010). We infer a twodimensional latent space and show that the capture time used as an informative prior helps to disambiguate pseudotime from branching structure.

### 3.1 Comparison with the DeLorean model

We have applied our model on three different datasets from three different organisms which have been also used by Reid and Wernisch (2016). The results produced by our model are similar to the DeLorean model, but our model converges significantly faster. All the experiments have been carried out by using the same experimental setup, that is the same model structure and initial conditions.

Windram *et al*. (2012) examined the effects of *Botrytis cinera* infection on *Arabidopsis thaliana*. Among the 150 genes described by Windram *et al*. (2012), we have used 100 genes for the inference process. The remaining 50 genes were left out as held-out genes and used further to validate the model as in Reid and Wernisch (2016). Fig. 1 shows the comparison of our method to the De-Lorean model. Fig. 1(**a**) shows the best and average, over 20 different initialisations, Spearman correlation between the actual capture time and the estimated pseudotime as the number of inducing points is increased. Both the best and average correlation values show that our method has faster convergence for a smaller number of inducing points than the DeLorean method. Fig. 1(**b**) depicts the fitting time required by both models for different number of inducing points. As our model uses the VFE approximation with an exact bound, it converges an order of magnitude faster than the DeLorean model which requires a sampling process. The problem with the sampling approach is that it requires initial burn-in time to fit the model which makes the inference slower and therefore problematic for larger datasets.

**Figure 1:**
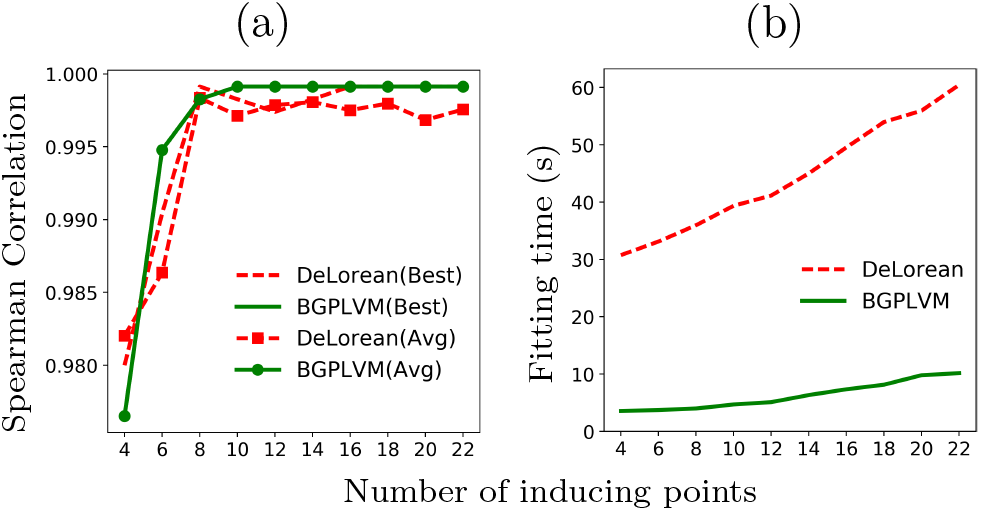
*Arabidopsis thaliana* microarray data (Windram *et al*., 2012): A comparison of performance and fitting time between the proposed method and DeLorean method. (**a**): Spearman correlation between the actual capture time and the estimated pseudotime for different number of inducing points. (**b**): Fitting time required by the models for the same experimental setups.

Reid and Wernisch (2016) defined the roughness statistic *R_g_* as the difference of consecutive expression measurements under the ordering given by pseudotime. Our model estimates smooth pseudotime trajectories which have close correspondence with the actual capture time points. To verify the smoothness of our predicted trajectory, we calculated the roughness statistics for the 50 held out genes. The average *R_g_* for all experiments in Fig. 1 is the same for both the DeLorean and Bayesian GPLVM approaches (0.71), reflecting the pseudotime similarity. For details, see supplementary.

Shalek *et al*. (2014) investigated the primary bone-marrow-derived dendritic cells of mouse in three different conditions. The time course data were collected using single-cell RNA-seq technology. They described several modules of genes which show different temporal expression patterns through the lipopolysaccharide stimulated (LPS) time course. Fig. 2(**a**) shows that our model correctly assigns two precocious cells to later pseudotime as in the DeLorean approach (see supplementary). Fig. 2(**b**) depicts the fitting time required by the both models for different number of inducing points and in all the cases the Bayesian GPLVM model converges significantly faster than the DeLorean model.

**Figure 2:**
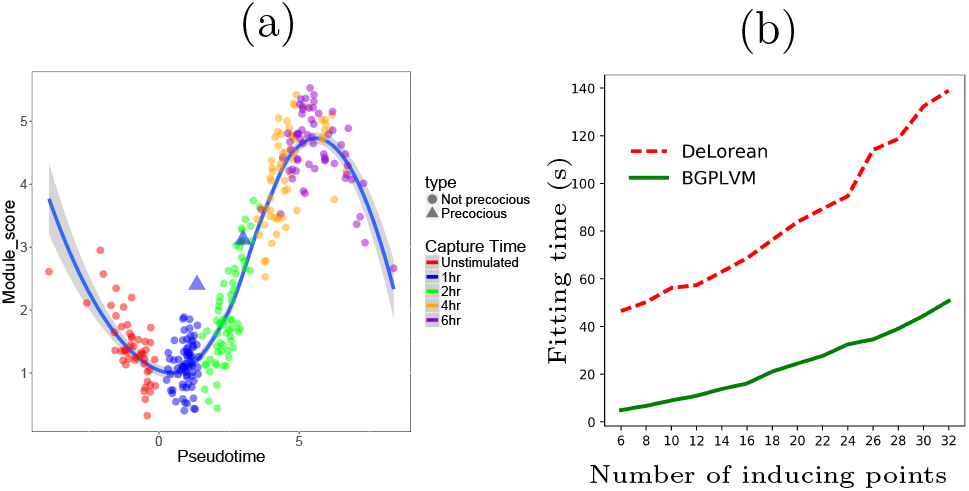
Mouse dendritic cells (Shalek *et al*., 2014): (**a**): The module score of core antiviral cells over pseudotime. The two precocious cells (plotted as triangles) have been placed in later pseudotimes than the other cells captured at 1 hour. A Loess curve (solid blue line) has been plotted thorough the data. (**b**): Comparison of fitting time required by both the DeLorean and our models for different number of inducing points while using the same experimental setups.

McDavid *et al*. (2014) examined the effect of cell cycle on single-cell gene expression across three human prostate cancer cell lines. To model the cyclic nature of the cell cycle, we have used a periodic kernel (Equation (6)). The DeLorean model requires 7h 31m to fit the model while our method uses 20 inducing points and takes only 4m 45s to converge whilst achieving similar error in recovering the cell cycle peak times (see supplementary). The DeLorean approach uses samples from 40 model initialisations to generate a full posterior GP whilst the BGPLVM only requires a single initialization as an analytic bound of the marginal likelihood is available. We also attempted to compare the fitting time required for different numbers of inducing points for this dataset but the sparse kernel used in the DeLorean packages results into non-invertible covariance matrices. Therefore the sparse approximation followed in the DeLorean package appears more fragile in cases of non-standard kernels such as the periodic kernel. The estimated pseudotimes are in good agreement with the cyclic behavior of the data. The model also predicts the cell cycle peak time of each gene with similar accuracy level of the DeLorean approach. See supplementary for the details of these results.

### 3.2 Scaling up the model to droplet-based single-cell data

To investigate the robustness and scalability of our method, we have applied it on droplet-based single-cell data. First, we have applied the model on single-cell RNA-seq data from mouse embryonic stem cells (ESC) generated using droplet barcoding (Klein *et al*., 2015). Klein *et al*. (2015) developed a method termed inDrop (indexing droplet) based on droplet microfluidics. They assayed the gene expression profiles and differentiation heterogeneity of mouse stem cells after leukaemia inhibitory factor (LIF) withdrawal. They captured the cells at *t* = 0, 2, 4 and 7 days and used their protocol to profile 2717 cells with 24175 observed transcripts. Haghverdi *et al*. (2016) have used this dataset for their analysis of diffusion pseudotime (DPT). They have applied their model on the cell cycle normalised data to infer DPT. We have used this cell cycle normalized data to assess the quality of the Bayesian GPLVM inferred pseudotime.

The inference process uses 2717 cells and 2047 genes. The model uses a RBF kernel (Equation (4)) to identify a smooth pseudotime trajectory. We have set the capture time prior variance to 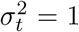. The pseudotime estimated by our model has a high rank correlation with both the actual capture time as well as the estimated pseudotime using DPT (Fig. 3).

**Figure 3:**
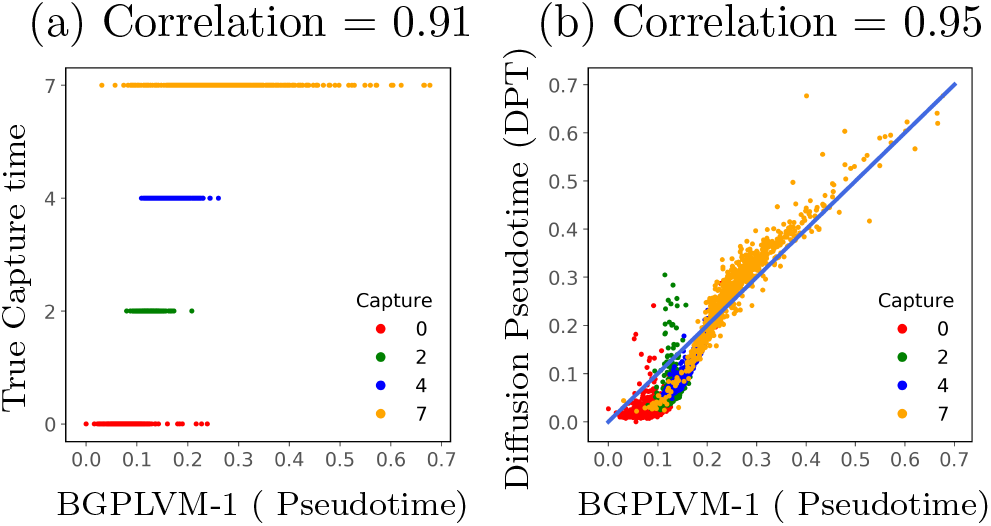
Mouse embryonic stem cells (Klein *et al*., 2015): Comparison of estimated pseudotime with the actual cell capture time and the pseudotime estimated using DPT. The points are coloured according to the actual cell capture times. The rank correlation is shown in the caption of each subplot.

As memory is a crucial resource when analysing large volumes of data, we also examine the effect of lower precision computations. We have examined the performance of our model under both 64 and 32 bits floating point precision. In both cases we observe a strong correlation with DPT (^~^0.94) but note a significant reduction in fitting time when using 32 bits precision. For 64 bits precision the algorithm take ^~^32 seconds to converge, whilst it takes only ^~^11 seconds to converge for 32 bits precision.

We also apply our method on a larger singlecell RNA-seq dataset to further demonstrate its scalability. Zheng *et al*. (2017) have presented a droplet-based technology that enables 3′ messenger RNA (mRNA) digital counting to encapsulate tens of thousands of single cells per sample. In their method, reverse transcription takes place within each droplet and barcoded complementary DNAs (cDNAs) have been amplified in bulk. The resulting libraries are then used for Illumina short-read sequencing. Their method has 50% cell capture efficiency and can process a maximum of 8 cells simultaneously in each run. Zheng *et al*. (2017) have assayed ^~^68k peripheral blood mononuclear cells (PBMCs) demonstrating the suitability of single-cell RNA-seq technology to characterise large immune cell populations.

We have applied our method using the top 1000 variably expressed genes ranked by their normalised dispersion (Zheng *et al*., 2017). We use a 2D GPLVM model with no capture time prior information and an RBF kernel (Equation (4)) with 60 inducing points. The inducing points and hyperparameters have been optimised jointly with model parameters and the algorithm takes ^~^10m to converge on a simple desktop machine^1^. To validate the GrandPrix result, we compare the clustering in the latent space with the clustering reported in Zheng *et al*. (2017). The latent space clustering is computed using the k-means algorithm with *k* = 10 clusters and we have used the adjusted rand index (ARI) (Hubert and Arabie, 1985) to evaluate it’s agreement with the cell labels reported in Zheng *et al*. (2017). The ARI has a value near to 0.0 if the cluster labelling is performed randomly and 1.0 for identical clusterings. A better solution is achieved when using t-SNE to initialise the latent space rather than PCA (see supplement), suggesting that it is worth considering different methods to initialise GrandPrix to improve the quality of the solution; a similar strategy is taken in Zwiessele and Lawrence (2016) where multiple dimension reduction methods are used to initialise a GPLVM model. We have also found that the GrandPrix ARI (0.54) is higher than the t-SNE method (0.51) showing an improvement over the initialisation used.

Further we have investigated the scalability of the model across varying number of CPU cores^2^. For simplicity only the 1-D latent positions are optimised, using fixed values for the kernel hyperparameters *l* = 1 and *σ*^2^ = 1 and the inducing points. In Fig. 4 we show the time required per iteration when using different number of CPU cores for both 32 and 64 bit precision. The computational benefits of lower precision are reduced as the number of cores is increased. We also note the diminishing returns of increasing the number of CPU cores; we see an approximately doubling of performance when increasing the number of cores from 2 to 4 but a reduced benefit when increasing from 8 to 16. We recommend a small number of cores is assigned to an individual model fitting, with any remaining resources assigned to perform multiple model fittings using different initial conditions. The latter is needed to alleviate the local minima problem inherent when fitting a Bayesian GPLVM model.

**Figure 4:**
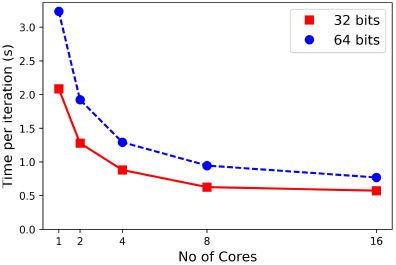
PBMCs with ^~^68k cells (Zheng *et al*., 2017): Time per iteration using 1, 2, 4, 8,16 CPU cores. The algorithm has been applied using both 32 and 64 bit floating point precision.

We can further increase the performance of the GrandPrix model by fixing rather than optimising the inducing point locations. This results in faster convergence without sacrificing accuracy given a sufficient number of inducing points is used (see supplementary). The effectiveness of this approach stems from the high amount of redundancy that is typical in larger datasets and offers a way to scale up the GrandPrix approach to datasets with a larger number of cells.

### 3.3 Extending the model to infer pseudotime-branching

To demonstrate the flexibility of our approach, we extend the model to 2-D latent spaces with a capture time prior on one latent dimension and apply it on single-cell qPCR data of early developmental stages in mouse (Guo *et al*., 2010). The gene expression profiles of 48 genes were measured across 437 cells. Cells differentiate from the single-cell stage into three different cell states in the 64 cell stage: trophectoderm (TE), epiblast (EPI), and primitive endoderm (PE).

Models with both informative and non-informative priors are examined. Both models use an RBF kernel (Equation (4)). Both models are initialized with identical values. For the informative prior, we set the capture time variance to 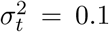. The informative prior (Fig. 5(**b**)) on capture time helps with the identifiability of the model as it aligns the first latent dimension (horizontal axis) with pseudotime and the second latent dimension (vertical axis) with the branching structure.

**Figure 5:**
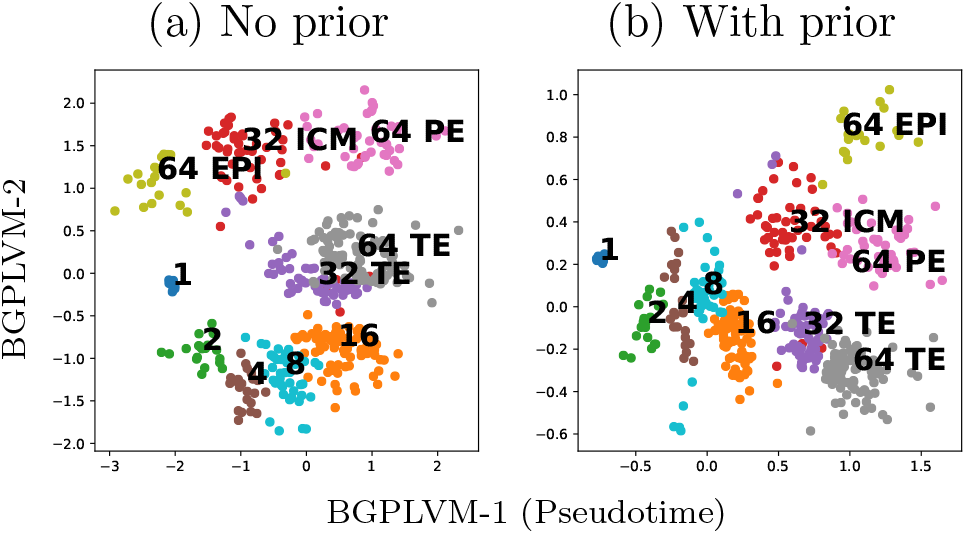
Single-cell qPCR of early developmental stages (Guo *et al*., 2010): Latent space reconstruction without and with prior. The bottom captures both developmental time and branching structure. The cell stage and type labels are also shown.

To investigate how the branching dynamics affect the estimation of pseudotime points, we have used our model to infer the 1-D pseudotimes with informative prior and compared it with the pseudotimes from the 2-D informative prior model (Fig. 6(a) and (b)). Both models were run from multiple initial conditions to ensure a good likelihood optimum was obtained. The 2-D model estimate of the pseudotime is found to have better correspondence with the actual capture time (correlation 0.84 vs 0.95), suggesting that the 1-D model is less able to align all variation with a pseudotime axis.

**Figure 6:**
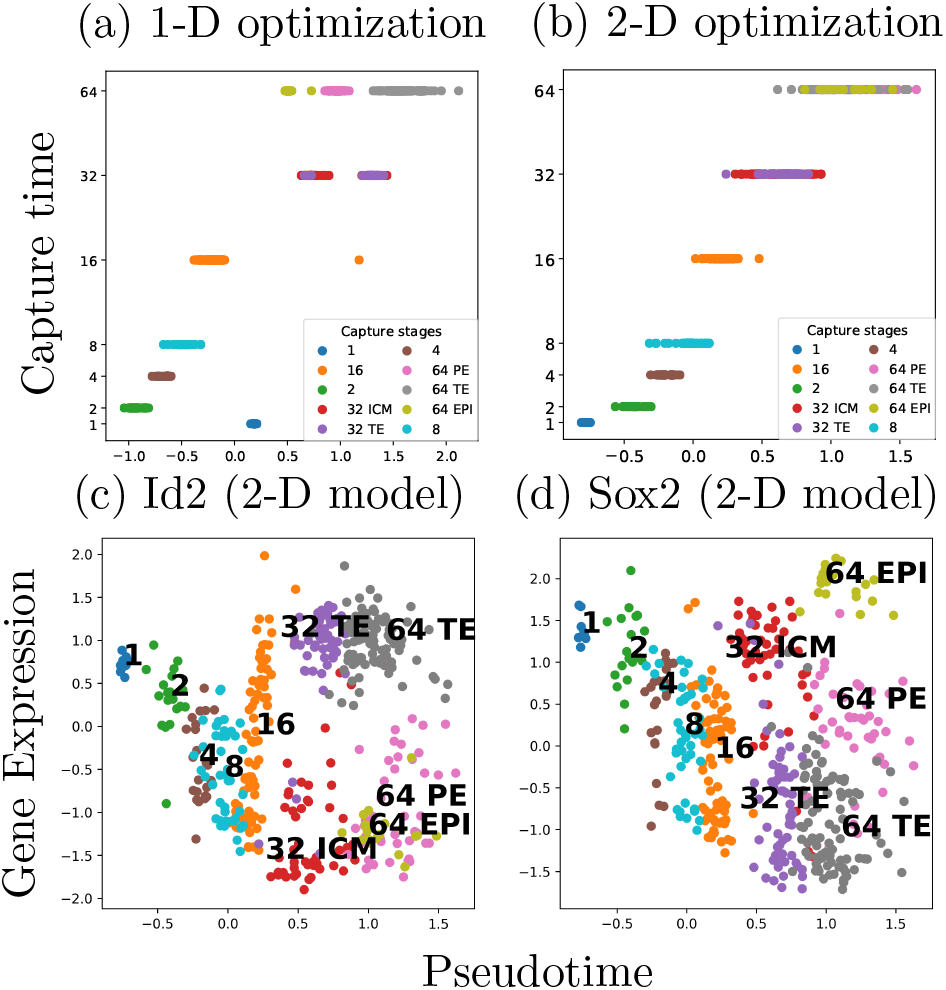
Single-cell qPCR of early developmental stages (Guo *et al*., 2010): (a) and (b): The actual capture times against the estimated pseudotimes from the 2-D and 1-D model with informative prior. (c) and (d): The expression profiles of the two known markers genes against the estimated pseudotime shows the time series experiments describing how the genes behaves differentially across the differentiation stages.

In Fig. 6(c) and (d), we have plotted the expression profiles of two marker genes against our estimated pseudotime points. *Id2* is a known marker gene for TE, thus it behaves differently in TE cells from the other two differentiation stages. It is differentially expressed between the stages TE and EPI, as well as between TE and PE. Similarly, Fig. 6 (d) shows that *Sox2* is differentially expressed between the stages TE and EPI, and between the stages PE and EPI. To see the expression profiles of the other differentially expressed genes across the differentiation stages, see the supplementary material.

## 4 Conclusion

Pseudotime estimation on single-cell genomics faces a number of challenges as the structure of the expression data is complex and non-linear. Many sources of variability, both biological and technical, introduce a significant amount of statistical uncertainty in the inference process. Here, we have used the Bayesian GPLVM model with informative priors to perform pseudotime estimation within a probabilistic framework. The model uses cell capture times as priors over pseudotime. Experimental results show that the properties of pseudotime ordering do not only depend on the data but also on the prior assumptions about the trajectory such as proximity to capture time, smoothness and periodicity.

The Bayesian GPLVM framework allows us to predict a number of latent dimensions along with associated uncertainty. A sampling-based Markov Chain Monte Carlo implementation of the Bayesian GPLVM is impractical for large number of cells because of its high computational complexity. We have developed our model on the basis of a sparse approximation that can generate a full posterior using only a small number of inducing points. Among a number of sparse approximation techniques, we have used the Variational Free Energy (VFE) approximation which has an exact bound to the marginal likelihood and avoids overfitting unlike the FITC approximation used by Reid and Wernisch (2016). To validate these claims, our approach has been tested on a variety of datasets from different organisms collected using different protocols. We find that our model has comparable accuracy to the DeLorean method for inferring the posterior mean pseudotime across all datasets used in Reid and Wernisch (2016) while converging considerably faster. The sources of the speed up are threefold: an analytic rather a numerically assessed variational bound, a more robust sparse approximation (VFE vs FITC) requiring fewer inducing points, and a scalable software implementation (Matthews *et al*., 2017) allowing for lower precision and GPU computation. The posterior mean from our model agrees closely with the posterior mean from DeLorean in all cases, but we find that the posterior variance of both the DeLorean and GrandPrix variational inference algorithms can be underestimated when compared to MCMC results (see supplementary Section 2.1). However, the DeLorean approach does not scale to datasets with more than a few hundred cells (Saelens *et al*., 2018). Our method therefore provides a practical approach to incorporate prior information into pseudotime estimation but at the cost of some loss in accuracy when assessing pseudotime uncertainties.

We have applied our model on droplet-based datasets to examine the robustness and scalability of our approach on much larger datasets. Our model successfully estimates pseudotimes for single-cell RNA-seq data of mouse embryonic stem cells (ESC) generated using the inDrop protocol. The Bayesian GPLVM estimated pseudotimes are in good agreement with DPT whilst providing all the benefits of a fully probabilistic model; namely quantification of uncertainty in the pseudotime estimation which has been shown to be of biological relevance (Campbell and Yau, 2016). To demonstrate our models scalability, we have measured its performance on a ^~^68k single-cell data of peripheral blood mononuclear cells and the model converges in 6 minutes on this large dataset.

Finally, we have applied the model on single-cell qPCR of early developmental stages to demonstrating its flexibility. We extended the model to higher dimensional latent spaces where the interaction of pseudotime with other factors, such as cell type differentiation, can be captured. We demonstrated the importance of this additional flexibility using a two-dimensional latent space where pseudotime is estimated jointly with the developmental branching structure. As extra latent dimensions can be used to describe other biological functions, the model can be extended to include additional prior information on the other latent dimensions; for example the prior could include information on branching dynamics extracted from the application of branching models such as Monocle (Qiu *et al*., 2017) and DPT (Haghverdi *et al*., 2016).

The model performs well across varying floating point precisions. For droplet-based datasets we have run the model using both 32 and 64 bit floating point precision and the algorithm produces similar estimation of pseudotime. We expect that in most cases, low precision will be sufficient to understand the behaviour of the system offering a way to further scale up our approach without the need for more expensive hardware. Mixed precision computations would also be possible with higher-precision computations performed only on the most numerically critical parts of the algorithm maintaining high accuracy whilst being significantly faster (Baboulin *et al*., 2009).

The analysis of single-cell data creates the opportunity to examine the temporal dynamics of complex biological processes where the generation of time course experiments is challenging or technically impossible. As single-cell data are becoming increasingly available in larger volumes, we believe scalable yet rigorous approaches such as the Bayesian GPLVM we have presented, will become ever more relevant. The flexibility of our approach can also reveal interesting biological facts such as identifying branching points in the differentiation pathways.

## Acknowledgements

We wish to thank John Reid for his help with the DeLorean code and data. We also thank James Hensman and all GPflow contributors for their help with the GPflow package. We express our gratitude to Jamie Soul and Robert Maidstone for their valuable insights.

## Footnotes

1 Intel(R) Core(TM) i5-3570 CPU @ 3.40GHz with 16 GB memory.

2 The hardware used was a 16-core Intel Ivy Bridge CPUs (E5-2650 v2, 2.60GHz) with 512 GB memory. TensorFlow version 1.0.0 and GPflow version 0.3.8.

## References

Baboulin, M., Buttari, A., Dongarra, J., Kurzak, J., Langou, J., Langou, J., Luszczek, P., and Tomov, S. (2009). Accelerating scientific computations with mixed precision algorithms. Computer Physics Communications, 180(12), 2526–2533.

Bauer, M., van der Wilk, M., and Rasmussen, C. E. (2016). Understanding probabilistic sparse gaussian process approximations. In Advances in Neural Information Processing Systems, pages 1533–1541.

Becher, B., Schlitzer, A., Chen, J., Mair, F., Sumatoh, H. R., Teng, K. W. W., Low, D., Ruedl, C., Riccardi-Castagnoli, P., Poidinger, M., et al. (2014). High-dimensional analysis of the murine myeloid cell system. Nature immunology, 15(12), 1181–1189.

Bendall, S. C., Davis, K. L., Amir, E.-a. D., Tadmor, M. D., Simonds, E. F., Chen, T. J., Shenfeld, D. K., Nolan, G. P., and Peer, D. (2014). Single-cell trajectory detection uncovers progression and regulatory coordination in human b cell development. Cell, 157(3), 714–725.

Buettner, F. and Theis, F. J. (2012). A novel approach for resolving differences in single-cell gene expression patterns from zygote to blastocyst. Bioinformatics, 28(18), i626–i632.

Buettner, F., Natarajan, K. N., Casale, F. P., Proserpio, V., Scialdone, A., Theis, F. J., Teichmann, S. A., Marioni, J. C., and Stegle, O. (2015). Computational analysis of cell-to-cell heterogeneity in single-cell rna-sequencing data reveals hidden subpopulations of cells. Nature biotechnology, 33(2), 155–160.

Campbell, K. and Yau, C. (2016). Order under uncertainty: robust differential expression analysis using probabilistic models for pseudotime inference. PLoS Computational Biology, 12(11).

Damianou, A. C., Titsias, M. K., and Lawrence, N. D. (2016). Variational inference for latent variables and uncertain inputs in gaussian processes. The Journal of Machine Learning Research, 17(1), 1425–1486.

GPy (since 2012). GPy: A gaussian process framework in python. http://github.com/SheffieldML/GPy.

Guo, G., Huss, M., Tong, G. Q., Wang, C., Sun, L. L., Clarke, N. D., and Robson, P. (2010). Resolution of cell fate decisions revealed by single-cell gene expression analysis from zygote to blastocyst. Developmental cell, 18(4), 675–685.

Haghverdi, L., Buettner, F., and Theis, F. J. (2015). Diffusion maps for high-dimensional single-cell analysis of differentiation data. Bioinformatics, 31(18), 2989–2998.

Haghverdi, L., Buettner, M., Wolf, F. A., Buettner, F., and Theis, F. J. (2016). Diffusion pseudotime robustly reconstructs lineage branching. Nature Methods, 13(10), 845–848.

Hubert, L. and Arabie, P. (1985). Comparing partitions. Journal of classification, 2(1), 193–218.

Ji, Z. and Ji, H. (2016). TSCAN: Pseudo-time reconstruction and evaluation in single-cell rna-seq analysis. Nucleic acids research, 44(13).

Klein, A. M., Mazutis, L., Akartuna, I., Tallapragada, N., Veres, A., Li, V., Peshkin, L., Weitz, D. A., and Kirschner, M. W. (2015). Droplet barcoding for single-cell transcriptomics applied to embryonic stem cells. Cell, 161(5), 1187–1201.

Lawrence, N. (2005). Probabilistic non-linear principal component analysis with gaussian process latent variable models. Journal of Machine Learning Research, 6(Nov), 1783–1816.

Lönnberg, T., Svensson, V., James, K. R., Fernandez-Ruiz, D., Sebina, I., Montandon, R., Soon, M. S., Fogg, L. G., Nair, A. S., Liligeto, U., et al. (2017). Single-cell rna-seq and computational analysis using temporal mixture modelling resolves th1/tfh fate bifurcation in malaria. Science immunology, 2(9).

MacKay, D. J. (1998). Introduction to gaussian processes. NATO ASI Series F Computer and Systems Sciences, 168, 133–166.

Marco, E., Karp, R. L., Guo, G., Robson, P., Hart, A. H., Trippa, L., and Yuan, G.-C. (2014). Bifurcation analysis of single-cell gene expression data reveals epigenetic landscape. Proceedings of the National Academy of Sciences, 111(52), E5643–E5650.

Matthews, A. G. d. G., van der Wilk, M., Nickson, T., Fujii, K., Boukouvalas, A., León-Villagrá, P., Ghahramani, Z., and Hensman, J. (2017). GPflow: A Gaussian process library using TensorFlow. Journal of Machine Learning Research, 18(40), 1–6.

McDavid, A., Dennis, L., Danaher, P., Finak, G., Krouse, M., Wang, A., Webster, P., Beechem, J., and Gottardo, R. (2014). Modeling bi-modality improves characterization of cell cycle on gene expression in single cells. PLoS Comput Biol, 10(7), e1003696.

Qiu, X., Mao, Q., Tang, Y., Wang, L., Chawla, R., Pliner, H. A., and Trapnell, C. (2017). Reversed graph embedding resolves complex single-cell trajectories. Nature methods, 14(10), 979.

Quiñonero-Candela, J. and Rasmussen, C. E. (2005). A unifying view of sparse approximate gaussian process regression. Journal of Machine Learning Research, 6(Dec), 1939–1959.

Rasmussen, C. and Williams, C. (2006). Gaussian processes for machine learning. MIT Press, Cambridge, Mass.

Reid, J. E. and Wernisch, L. (2016). Pseudotime estimation: decon-founding single cell time series. Bioinformatics, 32(19), 2973–2980.

Saelens, W., Cannoodt, R., Todorov, H., and Saeys, Y. (2018). A comparison of single-cell trajectory inference methods: towards more accurate and robust tools. bioRxiv, page 276907.

Shalek, A. K., Satija, R., Shuga, J., Trombetta, J. J., Gennert, D., Lu, D., Chen, P., Gertner, R. S., Gaublomme, J. T., Yosef, N., et al. (2014). Single-cell rna-seq reveals dynamic paracrine control of cellular variation. Nature, 510(7505), 363–369.

Shin, J., Berg, D. A., Zhu, Y., Shin, J. Y., Song, J., Bonaguidi, M. A., Enikolopov, G., Nauen, D. W., Christian, K. M., Ming, G.-l., et al. (2015). Single-cell rna-seq with waterfall reveals molecular cascades underlying adult neurogenesis. Cell Stem Cell, 17(3), 360–372.

Snelson, E. and Ghahramani, Z. (2006). Sparse gaussian processes using pseudo-inputs. In Advances in neural information processing systems, pages 1257–1264.

Titsias, M. K. (2009). Variational learning of inducing variables in sparse gaussian processes. In International Conference on Artificial Intelligence and Statistics, pages 567–574.

Titsias, M. K. and Lawrence, N. D. (2010). Bayesian gaussian process latent variable model. In International Conference on Artificial Intelligence and Statistics, pages 844–851.

Trapnell, C., Cacchiarelli, D., Grimsby, J., Pokharel, P., Li, S., Morse, M., Lennon, N. J., Livak, K. J., Mikkelsen, T. S., and Rinn, J. L. (2014). The dynamics and regulators of cell fate decisions are revealed by pseudotemporal ordering of single cells. Nature biotechnology, 32(4), 381–386.

Windram, O., Madhou, P., McHattie, S., Hill, C., Hickman, R., Cooke, E., Jenkins, D. J., Penfold, C. A., Baxter, L., Breeze, E., et al. (2012). Arabidopsis defense against botrytis cinerea: chronology and regulation deciphered by high-resolution temporal transcriptomic analysis. The Plant Cell, 24(9), 3530–3557.

Zheng, G. X., Terry, J. M., Belgrader, P., Ryvkin, P., Bent, Z. W., Wilson, R., Ziraldo, S. B., Wheeler, T. D., McDermott, G. P., Zhu, J., et al. (2017). Massively parallel digital transcriptional profiling of single cells. Nature communications, 8, 14049.

Zwiessele, M. and Lawrence, N. D. (2016). Topslam: Waddington landscape recovery for single cell experiments. bioRxiv.

